# The relationship between genomic variation and genetic load: insights from small island populations

**DOI:** 10.64898/2026.03.06.710193

**Authors:** Maëva Gabrielli, Andrea Benazzo, Roberto Biello, Alessio Iannucci, Daniele Salvi, Gentile Francesco Ficetola, Claudio Ciofi, Emiliano Trucchi, Giorgio Bertorelle

## Abstract

Small populations face high extinction risks. This can be explained by several non-genetic and genetic factors, the latter including the loss of genetic diversity and evolutionary potential, as well as the accumulation of harmful mutations (genetic load). Using whole-genome data from island populations with different effective sizes, we estimated genetic variation and load and explored the relationship between these quantities. An extremely small population of the Aeolian wall lizard, *Podarcis raffonei*, likely isolated for tens of thousands of years, shows the lowest genome-wide heterozygosity observed in wild eukaryotes (one polymorphic site every 300 kb on average). Despite this, its realized genetic load is comparable to that observed in another larger and more genetically variable population. Both populations have lower variation and higher load than the much more abundant sister species, the Sicilian wall lizard. These observations are consistent with the hypothesis that populations experiencing severe bottlenecks may persist for extended periods with extremely low genomic variation, provided that their burden of deleterious mutations remains within tolerable bounds.

## Introduction

The genetic factors contributing to the risk of extinction in a population are numerous and involve mutation, genetic drift, and inbreeding. Mutation frequently generates maladaptive variants (Muller, 1950), drift may subsequently increase their frequency, and mating among consanguineous individuals can expose deleterious recessive variants (Charlesworth and Charlesworth, 1999). Moreover, drift may also reduce beneficial genetic variation for future adaptation (Bijlsma and Loeschcke, 2012). Population size plays a crucial role in all these processes.

The partitioning of genetic load into components affecting current (realized load) and future (masked load) generations is crucial for understanding and potentially predicting extinction risk dynamics (Bertorelle et al., 2022). In large populations, individuals tend to accumulate more deleterious variants, defined as mutations with negative selection coefficients that reduce fitness, than individuals in small populations. This is due to a higher value of the parameter *Nμ* (effective population size multiplied by the mutation rate), as well as the fact that most mutations are mildly deleterious and recessive. Consequently, large populations exhibit a higher genetic load, but its impact is buffered as deleterious mutations in heterozygosis (the masked load) prevail over those in homozygosis (the realized load) (*e.g.* Humble et al., 2023; Smeds and Ellegren, 2023). Decreasing population size escalates drift effects, increasing both the frequency of deleterious variants, possibly up to fixation (genetic meltdown), and the level of inbreeding that may translate into unfit offspring (Charlesworth and Charlesworth, 1999). Only the removal of deleterious pairs of alleles in inbred offspring (genetic purging) can reduce the conversion of masked load into realized load and mitigate the subsequent fitness reduction and demographic decline (Crow, 1970).

Theoretical and simulation studies have explored the interplay between genetic variation and genetic load components to predict extinction risks under different demographic scenarios (Bertorelle et al., 2022). Whole-genome sequences have provided empirical insights by estimating genetic variation and load in single individuals (*e.g.* Humble et al., 2023; Pečnerová et al., 2023; Smeds and Ellegren, 2023). However, the effectiveness of whole-genome sequences as proxies for fitness in current and future generations, in particular for comparing populations and species with different population sizes and demographic histories, remains unclear. While some studies indicate the accumulation of realized genetic load and increased extinction risks as the main and most worrisome process in declining species (Dussex et al., 2021; Kleinman-Ruiz et al., 2022; Robinson et al., 2016; Von Seth et al., 2021), others suggest that purging is a relevant mechanism reducing vulnerability to extinction, particularly in populations experiencing prolonged periods of small population size (Ficetola et al., 2011; Grossen et al., 2020; Khan et al., 2021; Kleinman-Ruiz et al., 2022; Mathur et al., 2023b; Ochoa and Gibbs, 2021). Additionally, conflicting findings on the purging effect of highly deleterious mutations and the simultaneous accumulation of mildly deleterious alleles in small populations further complicate the understanding of fitness outcomes (Dussex et al., 2023; Grossen et al., 2020; Khan et al., 2021). Finally, the use of various load proxies and approaches to predict deleteriousness, coupled with the only recent appreciation of load partition into the realized and masked components, has led to inconclusive findings regarding extinction risk and general patterns of accumulation and expression of deleterious variants in natural systems (Bertorelle et al., 2022).

Here we investigate the interplay between population size, genomic variation, and genetic load in three insular lizard populations: two populations of the critically endangered (Corti et al., 2009a) Aeolian wall lizard (*Podarcis raffonei*) with census population sizes in the tens and hundreds of individuals, respectively, and, as a control, the sister species, the Sicilian wall lizard (*Podarcis waglerianus*), which is widely distributed across Sicily and satellite islands with a very large census population size (Corti et al., 2009b). The Aeolian wall lizard occurs in extremely small and isolated populations in the Aeolian archipelago (Fig. 1*A*) in southern Italy, with a population size ranging from 100 to 1,400 individuals across three islets and a small peninsula within an island, for a total population of about 2,000 individuals occupying an area of approximately five to six thousand square meters (Ficetola et al., 2021, 2018; Gippoliti et al., 2017; Salvi, 2023). Recent human disturbances and competition with the invasive Italian wall lizard (*Podarcis siculus*) have likely contributed to a reduction in the global population size and distribution range (Cascio and Corti, 2006).

**Figure 1.**
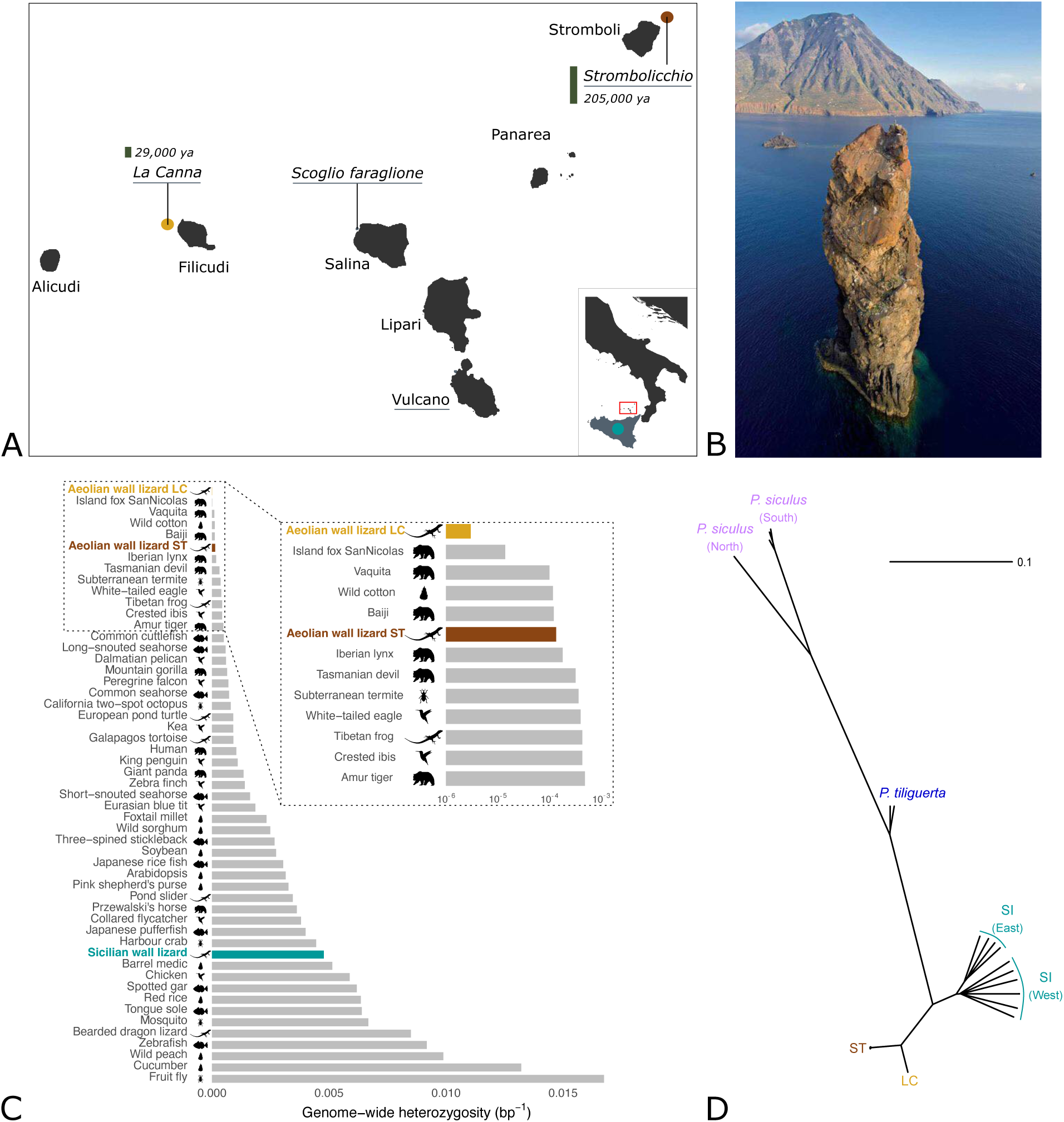
Distribution range, genetic diversity estimates and phylogenetic relationships of the Aeolian wall lizard. (A) Distribution range of the Aeolian wall lizard and the Sicilian wall lizard (bottom right, different scale). The red square in the inset indicates the position of the Aeolian archipelago. The Aeolian wall lizard is found in four islands and islets (underlined in grey), including our study sites of La Canna and Strombolicchio. The geologically estimated age of La Canna and Strombolicchio is indicated in italic. (B) Picture of the La Canna stack (credit: Piera Rapisarda), a volcanic neck 70m high and 1.6km distant from the island of Filicudi in the background. (C) Genome-wide observed heterozygosity in 53 eukaryotic species (Morin et al., 2021; Robinson et al., 2016), including the Aeolian wall lizard from La Canna (LC) and Strombolicchio (ST), and the Sicilian wall lizard. Silhouettes indicate the taxonomic group of the species included in this plot, namely plants, invertebrates, mammals, birds, fish, and reptiles and were retrieved from Phylopic (Gearty and Jones, 2023). (D) Neighbor-joining phylogenetic tree based on pairwise genetic distances between individuals, for 5 different population and species: the Aeolian wall lizard from La Canna (LC) and Strombolicchio (ST), the Sicilian wall lizard (SI), the Tyrrhenian wall lizard (*Podarcis tiliguerta*), and the Italian wall lizard (*Podarcis siculus*).

We focus on the populations of Strombolicchio (census population size of 500-700 individuals; Capula and Lo Cascio, 2011; Ficetola et al., 2021) and La Canna (census population size of about 100 individuals; Salvi, 2023; Fig. 1*B*). These are two volcanic islets located at approximately 1.5 km from the closest islands of Stromboli and Filicudi, respectively, where the Aeolian wall lizard is not found. The bathymetric profile suggests that the isolation of Strombolicchio from Stromboli and La Canna from Filicudi dates back to 7 to 10 kya (Cascio and Corti, 2006). Consequently, gene flow with the respective main islands might have ceased at around that time. It is worth noting that lizard populations might have been present in Strombolicchio much earlier than those of La Canna, as Strombolicchio was formed much earlier than La Canna (approximately 200 kya and 30 kya, respectively; Francalanci et al., 2013; Lucchi et al., 2013).

First, we characterized the genomic patterns of variation and divergence, and reconstructed the evolutionary history of two small populations of the Aeolian wall lizard and one large population of the Sicilian wall lizard. Then, we estimated and compared the genetic load across these three different groups. Despite an extreme level of inbreeding in the La Canna population, which behaves as a wild inbred strain, and distinct demographic histories, we found that the two populations of the Aeolian wall lizard exhibit a similar amount of realized genetic load. This confirm the expectation that the erosion of genetic variation is proportional to population size in isolated populations but also suggests that extinction can be avoided when genetic erosion is extreme but genetic load does not increase proportionally.

## Results

Mean individual sequencing depth was higher than 10x for all individuals except for two Italian wall lizard individuals, that were sequenced at a lower depth as they only act as outgroups. We obtained an average sequencing depth of 18.7x for the Aeolian wall lizard (22.5x and 15.3x for the La Canna and Strombolicchio individuals respectively) and 13.7x for the Sicilian wall lizard.

### 1) Genomic variation and population structure

The final dataset included 38 individuals (32 individuals sequenced in this study and six genomes from the literature; *SI Appendix*, Table S1) from four species: the Aeolian wall lizard (*P. raffone*i), the Sicilian wall lizard (*P. waglerianus*), the Italian wall lizard (*P. siculus*), and the Tyrrhenian wall lizard (*P. tiliguerta*). Thirty-nine millions of Single Nucleotide Polymorphisms (SNPs) were identified with a mean genotyping rate of 0.95: 10,810 biallelic SNPs were identified in La Canna, 335,876 in Strombolicchio, and 17 million in the Sicilian wall lizard (*SI Appendix*, Table S2). Single individuals confirmed this pattern, with Aeolian wall lizards showing much less heterozygous sites compared to Sicilian wall lizards, especially in La Canna (*SI Appendix*, Fig. S1). The average genome-wide heterozygosity was extremely low for both Aeolian wall lizard populations when compared to other eucaryotic species, with La Canna showing the lowest value documented so far (3.4x10^-6^, *i.e.* one SNP every 300 kb on average; Fig. 1*C*; *SI Appendix*, Fig. S2). Similar differences were found when considering only protein coding regions, with mean individual heterozygous counts of 94, 2,703 and 93,902 in La Canna, Strombolicchio and the Sicilian wall lizard, respectively. The extremely low genetic variation in La Canna and Strombolicchio was also observed in DNA regions where diversity is typically maintained by balancing selection. Specifically, one gene region from the class I and one from the class II Major Histocompatibility Complex (MHC) pathways, known to be highly variable in *Podarcis* lizards (Gaczorek et al., 2023), showed 22 and 13 polymorphic sites, respectively, in the Sicilian wall lizard, 6 and 0 polymorphic sites in Strombolicchio, and 0 polymorphic sites in both genes in La Canna.

The Principal Component Analysis revealed a clear separation between species and between La Canna and Strombolicchio individuals (*SI Appendix*, Fig. S3). Admixture analyses confirmed that species and populations were clearly separated, with no individuals showing mixed ancestry (*SI Appendix*, Fig. S4). Additionally, a phylogenetic reconstruction revealed that La Canna and Strombolicchio form two well-separated clades, characterized by shorter branches indicative of lower genetic diversity compared to the other species (Fig. 1*D*).

### 2) Inbreeding estimation and demographic history

The proportion of the genome in Runs of Homozygosity (F_ROH_) was about 0.98 in La Canna and 0.48 in Strombolicchio (Fig. 2*A*), suggesting extreme and high inbreeding levels in the two islets, respectively. La Canna and Strombolicchio populations had very different ROH length distributions (Fig. 2*B*), with an average ROH length of 50 Mb and 1.5 Mb in La Canna and Strombolicchio, respectively, while the Sicilian wall lizard was characterized by less than 100 regions in ROH that were always shorter than 300 kb and a F_ROH_ close to 0 (*SI Appendix*, Fig. S5)

**Figure 2.**
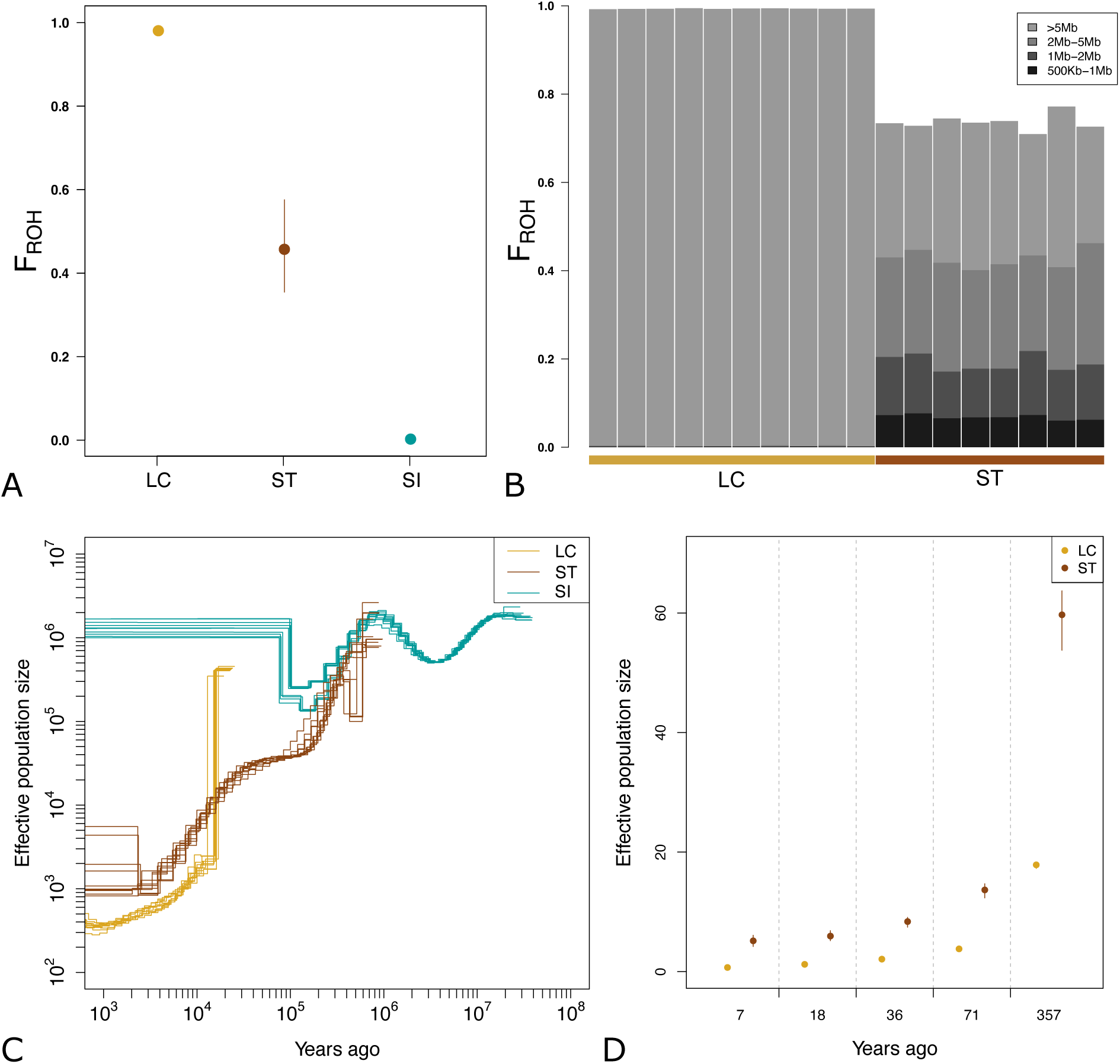
Inbreeding estimation and demographic history from effective population size variation though time. (A) F_ROH_ measured as the proportion of the genome in ROH longer than 1 Mb in the Aeolian wall lizard from La Canna (LC) and Strombolicchio (ST), and in the Sicilian wall lizard (SI). (B) Percentage of the genome in different lengths of ROH fragments for each individual of the La Canna (LC) and Strombolicchio (ST) populations. (C) Ancient *N_e_* variation through time as inferred from MSMC analyses, for the Aeolian wall lizard from La Canna (LC) and Strombolicchio (ST) and for the Sicilian wall lizard (SI). (D) Recent *N_e_* estimations from the proportion of the genome in different length of ROH fragments, for the Aeolian wall lizard from La Canna (LC) and Strombolicchio (ST).

The population size dynamics inferred by MSMC indicated a large population size for the Sicilian wall lizard (with some oscillation in the ancient past) and a constant demographic decline in both Aeolian lizard populations, particularly in La Canna (Fig. 2*C*). Around 200 kya, after a shared demographic decline, the trajectories of the Sicilian and the Aeolian wall lizard started diverging: the decline ended and likely reversed in the Sicilian wall lizard, with population sizes at high values (10^5^ – 10^6^), and the Aeolian wall lizard in Strombolicchio continued to decline gradually until 2 kya. Sometime earlier, around 20 kya, the shallow coalescent tree for the Aeolian wall lizard in La Canna allows the MSMC method to start the inference of the demographic dynamics of the individuals sampled on this rock: after an initial and intense drop, possibly due to a methodological artefact (Li and Durbin, 2011), the population size decreased more gradually, paralleling the dynamics of Strombolicchio, but with smaller absolute values. The orders of magnitude of the effective population sizes one thousand years ago are estimated at 10^2^, 10^3^ and 10^6^ in La Canna, Strombolicchio, and the Sicilian wall lizard, respectively. Interpretation of these absolute numbers requires great caution (*e.g.* the mutation rate is approximate), but they correspond to the estimated relative population sizes of these three groups of individuals. Recent effective population sizes in La Canna and Strombolicchio estimated from ROH patterns suggested even smaller population sizes in the last 500 years, with an initial decline to a modern effective population size of very few individuals (Fig. 2*D*, *SI Appendix*, Table S3). The divergence between the populations on La Canna and Strombolicchio was estimated using the SMC++ approach to have occurred approximately 14,000 years ago. However, given that both La Canna and Strombolicchio were likely colonized from the larger and geographically distant islands of Filicudi and Stromboli, respectively (see Fig. 1*A*), where the species is now extinct, this divergence time should not be interpreted as the time of colonization or separation of these two small islets. Rather, it provides a temporal framework for understanding the broader colonization history of the archipelago.

### 3) Genetic load and purging

Genetic load was investigated using two distinct approaches based on genomic annotations (SnpEff) and evolutionary conservation (GERP), at the individual and population levels. At the individual level, assuming that most deleterious mutations are recessive, homozygous and heterozygous genotype counts of derived deleterious mutations can be used as indicators of the realized genetic load (expressed and reducing the fitness of an individual) and the masked genetic load (unexpressed and not affecting individual fitness), respectively. The Aeolian and the Sicilian wall lizards showed opposite patterns at the genetic load partitions (*i.e.*, masked and realized), consistent with their different population sizes. Regardless of the approach used to identify deleterious mutations (missense, nonsense, high-GERP-score, or combination of these criteria), the Aeolian wall lizard showed approximately 30 to 900 times less masked load than the Sicilian wall lizard, and both Aeolian wall lizard populations showed approximately twice the realized load observed in the Sicilian wall lizard (Fig. 3 and *SI Appendix*, Tables S4 and S5). The masked load was much lower in La Canna than in Strombolicchio (Wilcoxon test, p-value=1.22e^-4^, 1.16e^-4^ and 1.24e^-4^ for missense, nonsense and high-GERP-score mutations, respectively), reflecting the different levels of genomic variation. Conversely, the realized load did not significantly differ between the two populations (p-value=0.22, 0.27 and 0.27 for missense, nonsense and high-GERP-score mutations, respectively).

**Figure 3.**
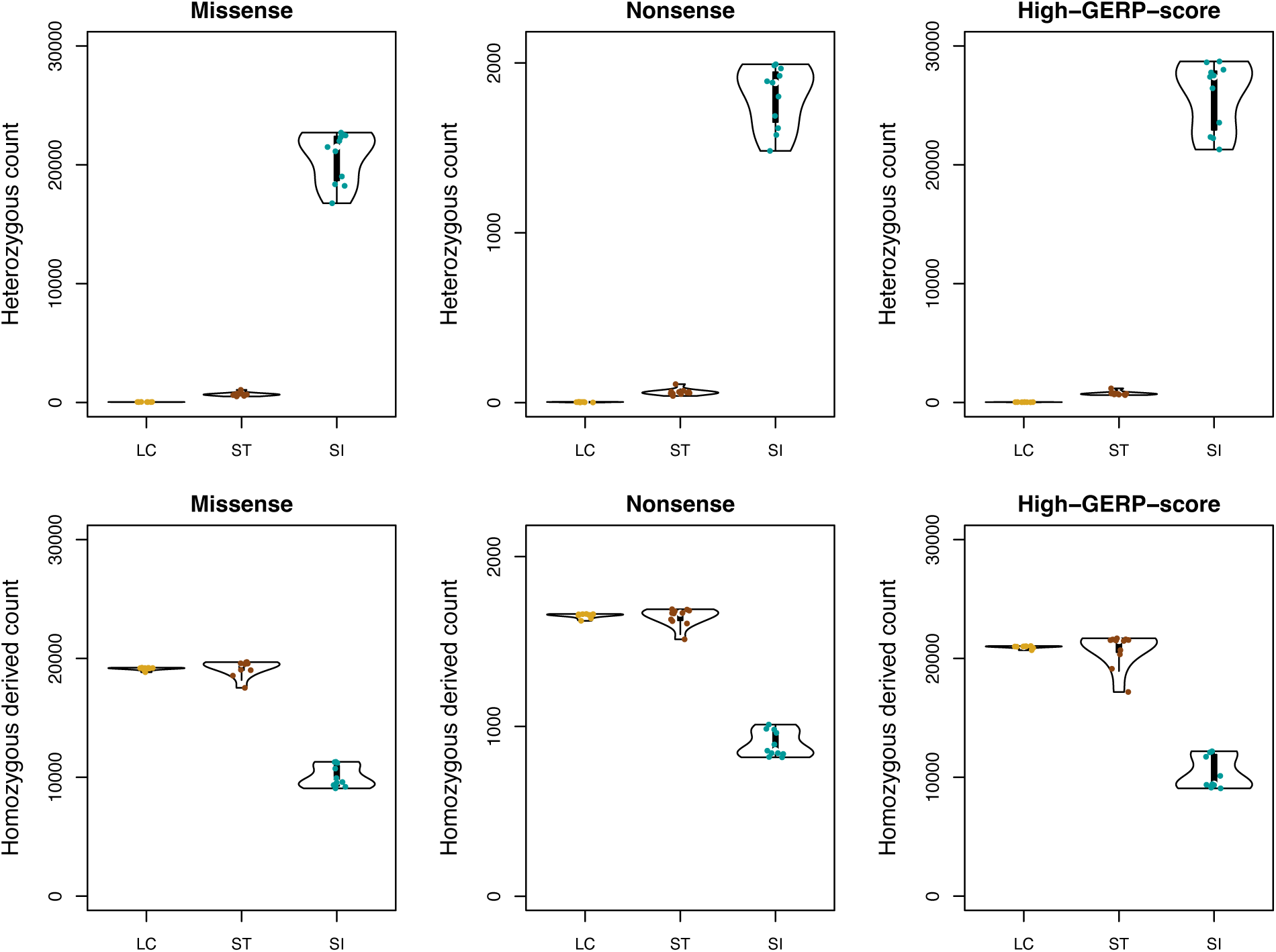
Genetic load estimations for the Aeolian wall lizard from La Canna (LC) and Strombolicchio (ST), and in the Sicilian wall lizard (SI). Number of genotypes for different classes of putatively deleterious mutations: missense (slightly deleterious mutations) and nonsense (highly deleterious mutations), classified using SnpEff, and high-GERP-score mutations (highly deleterious mutations), estimated from GERP (GERP score>4). Upper panels are heterozygous counts (proxies for the masked genetic load), and lower panels are homozygous derived counts (proxies for the realized genetic load).

To estimate the genetic load at the population level, thus considering how much individuals tend to share deleterious mutations, we computed the number of deleterious mutations at high frequency within the population (derived allele frequency higher than 0.9), following Mathur et al. (Mathur et al., 2023a). The population genetic load was higher in the Aeolian wall lizard than in the Sicilian wall lizard (5 to 6 times higher depending on the class of mutation considered), and very similar in the La Canna and Strombolicchio populations (*SI Appendix*, Table S6).

Finally, we found some partial evidence of purging in the small populations of the Aeolian wall lizard when compared to the much larger population of the Sicilian wall lizard. First, the total number of deleterious alleles was higher in Sicilian wall lizards compared to Aeolian wall lizards for all classes of putatively deleterious mutations and was lower in La Canna than in Strombolicchio (*SI Appendix*, Fig. S6). More explicitly, the number of missense, nonsense, and high-GERP-score mutations decreased respectively by 3.9%, 6.6% and 9.2% from the Sicilian wall lizard to the Aeolian wall lizard in Strombolicchio, and respectively by 5.9%, 7.9% and 9.8% from the Sicilian wall lizard to the Aeolian wall lizard in La Canna. Considering that the number of derived synonymous mutations in both Strombolicchio and La Canna was 5.0% and 4.4% lower than in the Sicilian wall lizard, respectively (this reduction is not expected theoretically under nucleotide independence, but has been observed in other studies (Kleinman-Ruiz et al., 2022) and it can be related to linkage effects), the higher reductions for the nonsense and high-GERP-score mutations can be interpreted as evidence of purging. Second, the difference in the total number of deleterious alleles between La Canna or Strombolicchio and the Sicilian wall lizard was lower for mildly deleterious mutations (missense mutations) compared to highly deleterious mutations (nonsense or high-GERP-score mutations; *SI Appendix*, Fig. S7, *SI Appendix*, Table S7). On the other hand, the expectation that under purging the Aeolian wall lizard should show a larger depletion of derived alleles with high GERP scores when compared to the Sicilian wall lizard is only partially met (SI Appendix, Fig. S8). The frequency distributions of GERP scores differed significantly among the three groups (pairwise two-sample Kolmogorov-Smirnov tests, p < 0.0001 for all comparisons; SI Appendix, Fig. S8), as well as the mean scores (Kruskal-Wallis test, p < 0.0001) and the proportion of variants with GERP values higher than 4 (chi-square test: χ2 = 78.393, d.f. = 2, P < 0.0001). The differences are however very weak: the average GERP score was 1.68, 1.69 and 1.70 in the Aeolian wall lizard from La Canna and Strombolicchio, and in the Sicilian wall lizard respectively, and the proportion of alleles with GERP scores larger than 4 increased from 4.96% to 5.04% to 5.24% in the same three groups. Statistically significant differences are clearly obtained considering that very large sample sizes are involved, but considering also the errors associated with the GERP score, the biological significance of these differences are probably minor.

## Discussion

The Aeolian and Sicilian wall lizard system appears as an extreme and ideal natural experiment to empirically explore the biological limits of genetic variation and load, and their relationship, in viable populations. The genomes of three groups of individuals were sequenced: Aeolian wall lizards from La Canna, Aeolian wall lizards from Strombolicchio, and Sicilian wall lizards. Assuming a genomic variation level of one in La Canna individuals, this value increases to 40 in Strombolicchio and 1,400 in Sicily. The estimated inbreeding coefficient was close to 1, 0.5, and 0 in La Canna, Strombolicchio, and Sicily, respectively. The reconstructed demographic trajectories of the lizard populations in Strombolicchio and La Canna are compatible with the estimated ages of these islands. Individual genomes from Strombolicchio indicate an ancient decline between 1 Mya and 200-300 kya, shared with the sister species (the Sicilian wall lizard) and possibly related to Pleistocene climatic events affecting the pre-colonization ancestors. Since then, the Aeolian wall lizard in Strombolicchio continued to decline, contrary to what happened to the Sicilian wall lizard. This decline is likely related to the erosion of genetic variation that occurred during and after the colonization of this island which emerged around 200 kya (Francalanci et al., 2013). The very low level of genomic variation observed in La Canna allows only to reconstruct the demographic trend for a more recent time window. Starting a few tens of thousands of years ago, the effective population size shows a rapid decline, compatible with a recent colonization after the emergence of this volcanic neck approximately 30 thousands of years ago (Lucchi et al., 2013). The extremely limited area of suitable habitat likely imposed a strong and persistent constraint on population size thereafter (Salvi, 2023). The complex history of divergence and colonization across the islands and islets is difficult to reconstruct, especially considering that the species, once likely widespread throughout the Aeolian archipelago (Paris et al., 2024), is now restricted to just three tiny islets (Scoglio Faraglione, Strombolicchio, and La Canna) and a narrow peninsula (Capo Grosso), and that only two of these populations were included in our study. However, the estimated divergence time of 14,000 years between La Canna and Strombolicchio supports a colonization followed by long-term isolation and strong genetic drift, leading to the observed genetic structure.

### The relationship between genetic variation and load

Recent empirical studies have gathered evidence for an increased genetic load in small and endangered populations across various species (*e.g.* Humble et al., 2023; Quinn et al., 2024). Small populations, usually characterized by low genetic variation and high inbreeding levels, tend to be characterized by an elevated realized load, whereas larger populations may experience an elevated masked genetic load (Humble et al., 2023; Smeds and Ellegren, 2023). Our results partially support this growing body of evidence as far as the two sister species of lizard (the Aeolian wall lizard and the Sicilian wall lizard) are compared. However, when comparing the two Aeolian wall lizard populations, the genetic load was not inversely correlated to the population size and the level of genomic variation, and was not directly related to the inbreeding coefficient. Despite their vastly different inbreeding levels and effective population sizes, individuals from La Canna and Strombolicchio exhibit, on average, similar numbers of deleterious homozygous sites and similar numbers of frequent deleterious mutations. The extent of genetic load is shaped by multiple aspects of demographic history, including inbreeding dynamics, the presence of ancestral masked load, and the potential for purging (Kyriazis et al., 2021; Pekkala et al., 2012). Still theoretical and simulation studies have shown that small populations accumulate realized load and fix deleterious mutations more readily, largely due to reduced efficacy of selection (Bertorelle et al., 2022; Dussex et al., 2023). Our empirical data from La Canna and Strombolicchio do not fully align with this expectation andsuggest that the fraction of genetic load directly affecting individual fitness (the realized genetic load) may approach a plateau in these two islets, potentially reflecting an upper bound compatible with population persistence. One possible explanation is that individuals in Strombolicchio and especially in La Canna already carry a high burden of homozygous deleterious mutations, such that additional increases in realized load may be poorly tolerated.

If this is true, we can speculate that both populations on Strombolicchio and La Canna are close to a mutational meltdown, even though their levels of variation differ, and that extinction occurred on other islands or rocks where the load threshold had been exceeded. Clearly, we do not propose a universal or absolute threshold. The levels of realized load observed in La Canna and Strombolicchio likely reflect context-specific tolerances shaped by the ecological simplicity and extreme isolation of these islets. In such settings, reduced predation, interspecific competition, or pathogen pressure may buffer the fitness impacts of some deleterious variants. It is therefore possible that similar or even lower levels of load could have more severe fitness consequences in more complex or stressful environments. This underscores the importance of integrating both genetic and ecological perspectives when interpreting genetic load in terms of extinction risk.

The existence of two small populations that differ by a factor of 40 in genomic diversity, with the smaller one being almost an inbred strain, yet exhibiting comparable realized load, raises a fundamental question: what are the relative roles of genetic variation and genetic load in the persistence or extinction of populations? Clearly, both factors are important: the correlations between genetic variation and adaptive potential, and between genetic load and reduced fitness, are well-supported both theoretically and empirically. However, if we extend our results to other systems, we should conclude that absence of variation can be better tolerated in small populations than increased genetic load, and historical extinctions due to genetic factors might have been driven more often by excessive generic load than by low genetic variation. The persistence of reproductively active populations with very low genomic diversity (Robinson et al., 2022, 2016), some evidence of a correlation between load and extinction (Rogers and Slatkin, 2017), and the weak correlation between genetic variation and IUCN Red List categories (Schmidt et al., 2023) are compatible with this idea, but further empirical, theoretical, and simulation-based testing of this hypothesis across different biological systems is required.

Populations or species with low or very low levels of genomic variation may still persist due to the presence of a few key genes with substantial levels of variation. This has been observed for immune system and/or olfactory receptor genes in the Apennine brown bear (Benazzo et al., 2017) or the Channel island fox (Robinson et al., 2016). Genetic variation at the MHC regions we analyzed was absent in the La Canna population and very limited in Strombolicchio but was relatively high in the same genomic regions in the Sicilian wall lizard. This analysis, while acknowledging the challenges associated with using genomic data to characterize complex MHC loci, suggests that these populations may have persisted despite limited diversity at some of the genes typically among the most variable and subject to balancing selection in vertebrates. The isolation and tiny size of these islets potentially limit the arrival of new pathogens, thereby reducing selective pressure on immune-related genes and reducing the importance of maintaining variation at these loci.

### Challenges in estimating genetic load

Genomics predictions of mutation deleteriousness have large potential for the study and the management of endangered species (Bertorelle et al., 2022; Dussex et al., 2023; van Oosterhout, 2020). However, at least three types of warning should be considered: biological, bioinformatical, and statistical. Biologically, i) the dominance architecture of most mutations of interest is not known, and deleterious mutations are usually considered fully recessive (Bertorelle et al., 2022); ii) rare variants specific to a lineage may be adaptive and not necessarily deleterious, even if absent in large multispecies alignments (Pagel et al., 2017); and iii) predicted deleterious variants in genes with low expression rate may have minor effects on individual fitness (Trucchi et al., 2023). Bioinformatically, the number of deleterious alleles may depend on i) the filtering approach; ii) the sample depth of coverage; and iii) the distance between the focal species and outgroups, the latter of which affects the total number of derived alleles. Statistically, i) reliable selection coefficients for each mutation are very difficult to obtain and their proxies such as GERP scores are affected by errors (Huber et al., 2020); and (ii) false positives (i.e., neutral variants classified as deleterious) may confound the pattern. Regarding false positives, a well-known issue when thousands of SNPs are analyzed, empirical studies have shown that GERP scores correlate with experimentally measured fitness effects in humans and dogs, supporting their general utility for identifying deleterious variants (Henn et al., 2015; Marsden et al., 2016). It is however possible that in case of strong purging effects that drastically reduce the number of true positives, differences or similarity between the load estimated in different populations may be only affected by the number of false positives.

More standardized load metrics, and functional studies to validate *in vivo* or *in vitro* the negative impact of variants bioinformatically estimated, are needed to increase the reliability of load estimates. Also, future work incorporating simulations or experimental validation will be essential to better understand the impact of false positives under different demographic and selective scenarios. All these improvements in estimating genetic load from genomic data will be crucial to test our hypothesis regarding the existence of a maximum load threshold, whose absolute value likely depends on the species’ ecology as well as its demographic and evolutionary history, and which may influence population persistence more directly than overall levels of genetic variation.

### Practical implications

If supported by additional analyses and broader datasets, the results of our study could have important implications for the conservation of endangered species. Preventing genetic erosion to enable adaptation to changing environments must always be pursued, but conservation strategies should also prioritize the protection of populations or species with intermediate levels of variation yet substantial realized load. Focusing on the Aeolian wall lizard populations, conservation efforts should be equally prioritized for La Canna and Strombolicchio, as both groups appear to be approaching a potential load threshold at which any additional genetic or environmental pressure could disproportionately increase extinction risk. The survival of this species depends of course also on the other two known populations in the archipelago, where genomes should be urgently sequenced. However, the wild and almost clonal population of La Canna should be strictly protected and closely monitored to better understand the relationship between genomic patterns, phenotypes, and extinction risks.

## Methods

### Population sampling and genomics data generation

A total of 32 lizards were sampled in the Aeolian archipelago and the island of Sicily. Ten individuals of Aeolian wall lizards were sampled on the stack of La Canna (38°34’56.13”N – 14°31’16.61”E; Fig. 1*B*), in a small terrace at 50 m above sea level on the eastern slope of the stack. Ten individuals were sampled on the islet of Strombolicchio (38°49’01.8"N - 15°15’06.7"E). Ten individuals of Sicilian wall lizards were sampled in Sicily, four individuals from the South-east and six individuals from the West of the island (see *SI Appendix*, Table S1 for details). Finally, two individuals of Italian wall lizards were sampled in Sicily. For each individual, the tip of the tail was cut, flask frozen in liquid nitrogen and later stored at -80°C. Individuals were released after sampling. Whole DNA was extracted using a PureLink Genomic DNA Mini Kit (Invitrogen). DNA integrity was assessed by 1.5% agarose gel electrophoresis and DNA concentration was measured using a Qubit 4 fluorometer Broad Range Assay (Invitrogen). Genomic libraries were constructed using an Illumina DNA PCR-Free Prep Kit (Illumina) according to the manufacturer’s protocol. Target coverage was 10-15X for all samples. Libraries were sequenced paired-end on an Illumina NovaSeq 6000 System using a 300-cycle S2 Reagent Kit v1.5. Read quality was evaluated with FastQC v0.12.1 (Andrews, 2010), and output reports were examined visually.

### Read mapping

A high-quality reference genome of the Aeolian wall lizard (Gabrielli et al., 2023) was used for read mapping. In addition to the 32 individuals collected and sequenced for the study, we used six *Podarcis* genomes from Yang et al. (Yang et al., 2021): one Aeolian wall lizard from Strombolicchio, one Sicilian wall lizard from Sicily, two Italian wall lizards from Sicily and the Tuscan islands, respectively, and two Tyrrhenian wall lizards from Sardinia and Corsica, respectively, the last two species serving as outgroups for our subsequent analyses. Reads were mapped to the reference genome using BWA version 0.7.17 (Li and Durbin, 2009) with default parameters, with an Optical Density (OD) value of 100 for our samples as they were generated using patterned flow cells and 2,500 for the other individuals. PCR and optical duplicates were tagged using the MarkDuplicates tool in the picard toolkit version 2.24.1 (http://broadinstitute.github.io/picard/). In our study, a PCR-free library preparation kit was used so that reads tagged as PCR duplicates were further untagged using a custom bash script. A number of statistics were computed to ensure the quality of the mapping (*SI Appendix*, Table S8, Fig. S9).

### Variant calling and filtering

An independent SNP calling was performed for the Aeolian wall lizards from La Canna and Strombolicchio, the Sicilian wall lizards, the Italian wall lizards and the Tyrrhenian wall lizards. SNP calling was performed using GATK (McKenna et al., 2010) v4 following the best practice guides for short variant discovery that is composed of three steps: (i) calling variants per sample using HaplotypeCaller; (ii) consolidate GVCFs to improve scalability using GenomicsDBImport; (iii) joint-genotype all the per-sample GVCFs using GenotypeGVCFs. HaplotypeCaller was run independently in each scaffold of each individual, using the option EMIT_ALL_CONFIDENT_SITES to output all sites (including non-variant sites). The outputs were generated in condensed non-variant blocks. A minimum mapping quality of 20 was used. Finally, all scaffolds were merged together to produced one unique VCF file per dataset using bcftools merge (Danecek et al., 2021). From the raw VCF file produced after SNP calling, we first extracted SNPs and indels using GATK SelectVariant. We then filtered indels by quality (minimum quality of 60) and then filtered out low-quality SNPs that were located within 5bp of the high quality indels, with the following criteria: QUAL < 60; QD < 2.0; FS > 60.0; MQ < 40.0; MQRankSum < -20.0; ReadPosRankSum < - 8.0; DP < meandepth/3; DP > meandepth*2; GQ < 10, using GATK VariantFiltration. We removed repeat regions, regions of high heterozygosity and filtered out sites with more than 25% of missing data using VCFtools (Danecek et al., 2011) and bedtools intersect (Quinlan and Hall, 2010). We also extracted the invariant sites and filtered out low-quality sites with a RGQ < 10. Finally, we merged high-quality SNPs and invariant sites for each group with bcftools concat and merged the VCF files of all groups using bcftools merge. We removed the two sexual chromosomes (Z and W) and restricted the analyses to the 18 autosomes

### Genomic diversity and population structure

We first quantified genomic diversity in our three groups of interest, the Aeolian wall lizard from La Canna, the Aeolian wall lizard from Strombolicchio and the Sicilian wall lizard, computing the number of SNPs per population and species and the individual number of heterozygous sites. We also computed the mean observed heterozygosity per population and compared the obtained estimates to genomic estimates of observed heterozygosity in a range of eucaryotes (Morin et al., 2021; Robinson et al., 2016).

We investigated genomic diversity in the Major Histocompatibility Complex (MHC), a region typically characterized by high variability in vertebrates. To identify relevant loci, we used data from Gaczorek et al. (2023), which characterized MHC regions in *Podarci*s lizards inhabiting the Iberian Peninsula. We focused on two genes, corresponding to a class I and class II MHC second exons, both showing high variability in Gaczorek et al. (2023). By performing a BLAST search (Altschul et al., 1990) of ten randomly selected MHC I alleles deposited by Gaczorek et al. (2023) against the *P. raffonei* reference genome (accession GCF_027172205.1), we consistently retrieved 15 accession numbers, all of which corresponded to the "H-2 class I histocompatibility antigen Q9 alpha chain-like" protein sequence. After filtering out isoforms of the same transcript and sequences shorter than 200 bp, six very similar sequences were retained. One of them, randomly selected, was used as a query to extract the homologous region in the genomes of lizards from La Canna, Strombolicchio, and the Sicilian wall lizard and estimate the number of SNPs in the three groups. We applied the same approach for MHC II alleles. Five accession numbers were retrieved corresponding to the "H-2 class II histocompatibility antigen, E-S beta chain-like" protein sequence. After filtering for isoforms and fragment length, two very similar sequences were retained and one of them was used to estimate the genetic variation in La Canna, Strombolicchio, and in the Sicilian wall lizard.

In order to analyze population and species clustering and ensure the global quality of our dataset, we first performed a Principal Component Analysis using Plink (Purcell et al., 2007). We then analyzed the shared ancestry between individuals using Admixture (Alexander et al., 2009) using sites segregating in the Aeolian, Sicilian and Italian lizards and 50 kb apart in order to have independent markers (resulting in 30,497 SNPs). Finally, we performed a phylogenetic tree based on pairwise genetic distances between individuals using Plink.

### Inbreeding levels

To quantify the levels of inbreeding in our three populations of interest, we used ROHan, a probabilistic method that infers local rates of heterozygosity and delineates Runs of Homozygosity (ROH) (Renaud et al., 2019). We run the program with a rhomu value of 5e^-5^ and a window size of 1 Mb. We then considered a region to be a ROH when the P(ROH) was higher than 0.9, and computed the proportion of the genome in ROH higher than 1Mb, (FROH), which serves as a genomic estimate of the inbreeding coefficient F. We also used a window size of 100kb in order to estimate the proportion of the genome in different classes of ROH length, and here removed two individuals (ST03 and ST04) for which the chains did not reach convergence.

### Demographic inferences

The long-term variation in population size was first investigated using MSMC2 (Schiffels and Wang, 2020; Wang et al., 2020). This analysis was run on the 18 different autosomes using three masks. The first was a callable mask that was computed using GATK CallableLoci on the bam files. We defined callable sites as sites with a minimum base quality and mapping quality of 20 and covered between one third and three times the mean individual depth. The second was a mappability mask generated using GEM v1.4.3 (Derrien et al., 2012) to determine uniquely mapping positions in the reference genome of the Aeolian wall lizard. The third was a repeat negative mask to exclude repetitive regions extracted from the output of RepeatMasker previously run on the reference genome of the Aeolian wall lizard . We used a default time patterning ("4+25*2+4+6"), with an initial theta/rho ratio (-r parameter) of 5 and a maximum 2N0 coalescent time (-N parameter) of 15 (default values). The recombination rate was set as variable in the MSMC algorithm. We used a generation time of two years and a mutation rate of 1e^-9^ mutations per site and per year as estimated from *Podarcis* whole-genomes (Yang et al., 2021).

The recent demographic history was investigated from the distribution of the ROH. The calculation of the coalescent times of ROH were estimated as: 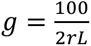 (Thompson, 2013), where g is the expected time back to the parental common ancestor when the IBD haplotypes forming an ROH coalesce, r is the recombination rate, and L is the length of the ROH in megabases. We used the recombination rate estimate of the chicken (*Gallus gallus*) of 2.8 cM.Mb^-1^, as done by Ochoa and Gibbs (Ochoa and Gibbs, 2021) , as this is, to our knowledge, the closest available estimate applicable to reptiles in the absence of species-specific data. We then derived the formula: 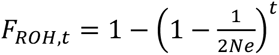 (Khan et al., 2021) to get an estimate of *N*_e_ from different time periods corresponding to different classes of ROH length.

To estimate the divergence time between the La Canna and Strombolicchio populations, we used SMC++ v1.15.4 (Terhorst et al., 2017), which implements a sequentially Markovian coalescent model to infer past population size changes and split times from unphased genomic data. As SMC++ treats missing data as homozygous reference genotypes, we first masked genomic regions with low or excessively high coverage, as well as those with insufficient base or mapping quality, using the same callable mask used for MSMC and computed using GATK CallableLoci. We generated .smc files for each autosome using the vcf2smc command, with the --mask option to exclude unreliable regions. We estimated marginal effective population sizes for each population using smc++ estimate, and applied the smc++ split command to fit a clean-split model between La Canna and Strombolicchio and infer the divergence time between the two populations. Analyses were run assuming the same mutation rate and generation time as in the MSMC analysis.

### Genetic load estimation

#### 1 - Ancestral state definition

A fundamental component of the genetic load estimation is the definition of the ancestral state. We therefore used two outgroups with different divergence times from the clade formed by the Aeolian and Sicilian wall lizard: the Tyrrhenian wall lizard from the Western islands group (divergence time: 16 My) and the Italian wall lizard (divergence time: 18 My) (Yang et al., 2021). We defined the ancestral allele as the allele present in at least three or the four Italian wall lizard individuals and the two Tyrrhenian wall lizard individuals using a custom python script. All sites where at least one of these outgroups was heterozygous were discarded to maximize the confidence in the ancestral allele definition. We then selected the sites for which at least one derived allele was present in the Aeolian and/or Sicilian wall lizard, but not fixed in these two species, resulting in 13,421,208 sites.

#### 2 – Annotation-based approach to estimate the genetic load

We first estimated the genetic load based on genomic annotations and classified mutations of different effects using SnpEff (Cingolani et al., 2012). Well-known functional effects of mutations in different genomic regions are used to classify a mutation. Mutations inside coding protein sequences (CDS) were categorized into three classes bases based on their predicted impact: LOW (synonymous); MODERATE (missense, *i.e.*, nonsynonymous), and HIGH (nonsense mutations including stop-gained, start-gained, start lost, splice acceptor, and splice donor variants). When one mutation was associated to multiple effects (due to different genes being affected by the same variant or one gene having multiple transcripts), the first one was considered, corresponding to the higher putative impact and to canonical over non-canonical transcripts.

#### 3 – Evolutionary conservation-based approach to estimate the genetic load

We then estimated the genetic load based on evolutionary conservation using GERP++ (Davydov et al., 2010). This approach builds on the idea that rare substitutions in genome phylogenies across species indicate strong constraints, so that variants in these positions are predicted to be deleterious. GERP scores were calculated on an alignment of 46 species including 16 reptiles (Roscito et al., 2022). As the tegu (*Salvator merianae*) was used as the reference genome for this later multi-species alignment, we first aligned the genome of the Aeolian wall lizard to the genome of the tegu (obtained in DNA Zoo, version 6; Roscito et al., 2018) using AnchorWave v1.0.1 (Song et al., 2022). We then used liftOver (UCSC; Kuhn et al., 2013) to change the coordinates of the tegu genome into the coordinates of the Aeolian wall lizard genome. Mutations with a GERP score higher than 4 were classified as putatively deleterious, following Smeds et al. (2023) and based on the distribution of the GERP scores of synonymous, missense and nonsense mutations from the genomic annotations (see *SI Appendix*, Fig. S10). Nonsense mutations had a higher GERP score (mean value: 2.3) than missense mutations (mean value: 1.9), that had a higher GERP score than synonymous mutations (mean value: -0.87; significant differences in mean scores from pairwise Wilcoxon tests; *SI Appendix*, Fig. S10), showing that inferences based on annotations and evolutionary conservation are congruent.

#### 4- Proxies of the genetic load components

Assuming that most deleterious mutations are recessive, homozygous and heterozygous genotype counts of derived deleterious mutations can be used as indicators of the genetic load components. For each class of putatively deleterious mutations, namely missense, nonsense and high-GERP-score mutations, the genetic load was first measured at the individual level and separated into two components: (i) the masked load (unexpressed and not affecting individual fitness) was estimated by the individual number of heterozygous genotypes and (ii) the realized load (expressed and reducing the fitness of an individual) was estimated by the individual number of homozygous genotypes for the derived allele (Bertorelle et al., 2022). We also computed these counts for high-GERP-score missense mutations and high-GERP score nonsense mutations. At the population level, we characterized genetic load as the number of deleterious alleles present in a population at high frequency (frequency of the derived allele higher than 0.9), following Mathur et al. (Mathur et al., 2023a). To ensure comparisons across populations were not confounded by patterns of neutral variation, we also compared the total number of derived alleles at synonymous SNPs.

#### 5 - Purging inference

To estimate the levels of purging in our system, we first contrasted the total number of deleterious alleles in the different populations. In case of purging, we expect a loss of deleterious alleles in small populations, as in Kleinmann-Ruiz et al. (Kleinman-Ruiz et al., 2022). We then computed the percentage of difference in the number of deleterious alleles between the Sicilian wall lizard and both populations of the Aeolian wall lizard for mildly deleterious mutations (missense mutations) and highly deleterious mutations (nonsense and high-GERP-score mutations). If purging is more efficient for highly deleterious mutations, the percentage of difference between the number of highly deleterious mutations between large and small populations should be higher than the one for mildly deleterious mutations (Grossen et al., 2020; Khan et al., 2021). Finally, we computed the average coefficient of deleteriousness of mutations based on GERP scores for the three groups of individuals. In case of purging, lower values of this average are expected in small populations where highly deleterious mutations have been eliminated (Dussex et al., 2021).

#### 6 – Statistical analyses

The two-tailed Wilcoxon test was used to compare means in the "Genetic load and purging" section. Considering the very low number of tests we performed, and the highly significant p-values obtained when the null hypothesis was rejected, no correction for multiple testing was applied. We compared the distributions of GERP scores between the three groups using Kruskal-Wallis and Kolmogorov-Smirnov tests and we compared the proportion of high-GERP-score mutations (mutations with GERP score larger than 4) using a Chi-square test.

## Acknowledgments

We thank the mountain guide Lorenzo Inzigneri for its help in the sampling in La Canna. We thank Silvia Fuselli for her important contribution in the analysis of MHC variation. Sampling authorization was given under Prot. 45382, Rif. 35644/2019. This work was supported by the University of Ferrara (Italy) and funded by the MIUR PRIN 2017 grant 201794ZXTL to G.B.

## Author Contributions

Designed research: E.T. and G.B.

Performed research: M.G., G.B., A.B., E.T., R.B., C.C., D.S., and A.I.

Analyzed data: M.G. and G.B.

Wrote the initial draft: M.G. Wrote the paper: M.G. and G.B

Interpreted the results: M.G., A.B., E.T., D.S. and G.B Discussed and approved the final manuscript: all the authors.

## Data availability

Raw sequencing data are available under NCBI BioProject PRJNA1089471.

